# Encoding Information in Synthetic Metabolomes

**DOI:** 10.1101/627745

**Authors:** Eamonn Kennedy, Christopher E. Arcadia, Joseph Geiser, Peter M. Weber, Christopher Rose, Brenda M. Rubenstein, Jacob K. Rosenstein

## Abstract

Biomolecular information systems offer numerous potential advantages over conventional semiconductor technologies. Downstream from DNA, the metabolome is an information-rich molecular system with diverse chemical dimensions which could be harnessed for information storage and processing. As a proof of principle of postgenomic data storage, here we demonstrate a workflow for representing abstract data in synthetic metabolomes. Our approach leverages robotic liquid handling for writing digital information into chemical mixtures, and mass spectrometry for extracting the data. We present several kilobyte-scale image datasets stored in synthetic metabolomes, which are decoded with accuracy exceeding 98-99% using multi-mass logistic regression. Cumulatively, >100,000 bits of digital image data was written into metabolomes. These early demonstrations provide insight into the benefits and limitations of postgenomic chemical information systems.

## Introduction

The metabolome is the complete set of small molecules found in a biological system [1]. The properties of this set of compounds are an amplified and dynamic measure of an organism’s genome, transcriptome, proteome, and environment [2]. This makes the metabolome an incredibly information-rich system, which displays diverse chemical, structural and biological dimensions [3–5]. Despite this complexity, improvements in protocols and efficient mass spectrometry (MS) have enabled metabolomic disease screening and drug discovery [6–12]. These technologies are supported by continually improving statistical tools and databases [13, 14]. As these tools advance, they may also suggest exciting alternative applications for metabolomics.

For inspiration, we observe that researchers have mimicked living systems by using DNA [15] for long-term archival information storage [16, 17], building on rapid advances in sequencing technology. Given recent progress in proteomic and metabolic profiling tools [18–21], it is timely to explore if the metabolome can also be used in a complementary way for information representations.

Whereas DNA and proteins are often large molecules which exist in small numbers, metabolites are higher in number, smaller in mass, and more structurally and energetically diverse. Like DNA, metabolites are biologically ubiquitous, and their primary pathways and processes are conserved across species [22]. The power of DNA as an information carrier comes from the combinatorial complexity that can exist within one polymer [23]. By contrast, the power of the metabolome is in the diversity of many co-existing molecules which can interact, or be acted upon, in complex combinations [5].

Non-genomic molecular data storage has also been demonstrated using fluorescent dyes on polymer films [24] and rotaxanes [25]. Other demonstrations have utilized collections of fluorophores which interact with information-bearing compounds in statistically identifiable ways [26]. However, all of these methods encode information into the state of a single compound at one time.

In this paper, we encode abstract binary data into the chemical composition of thousands of spatially arrayed nanoliter volumes (Fig. 1a). Each volume (‘spot’) contains a prescribed mixture from a library of purified metabolites – a synthetic metabolome. A key strength of this work is that any chemical library could function equivalently. Metabolites hold particular potential, because they provide access to well-regulated interconversion networks, materials, and databases which could facilitate computational operations on chemical data. The presence or absence of one metabolite in one spot encodes one bit of information. Therefore, the total number of bits stored in one spot is equal to the number of available library elements [27].

**Figure 1:**
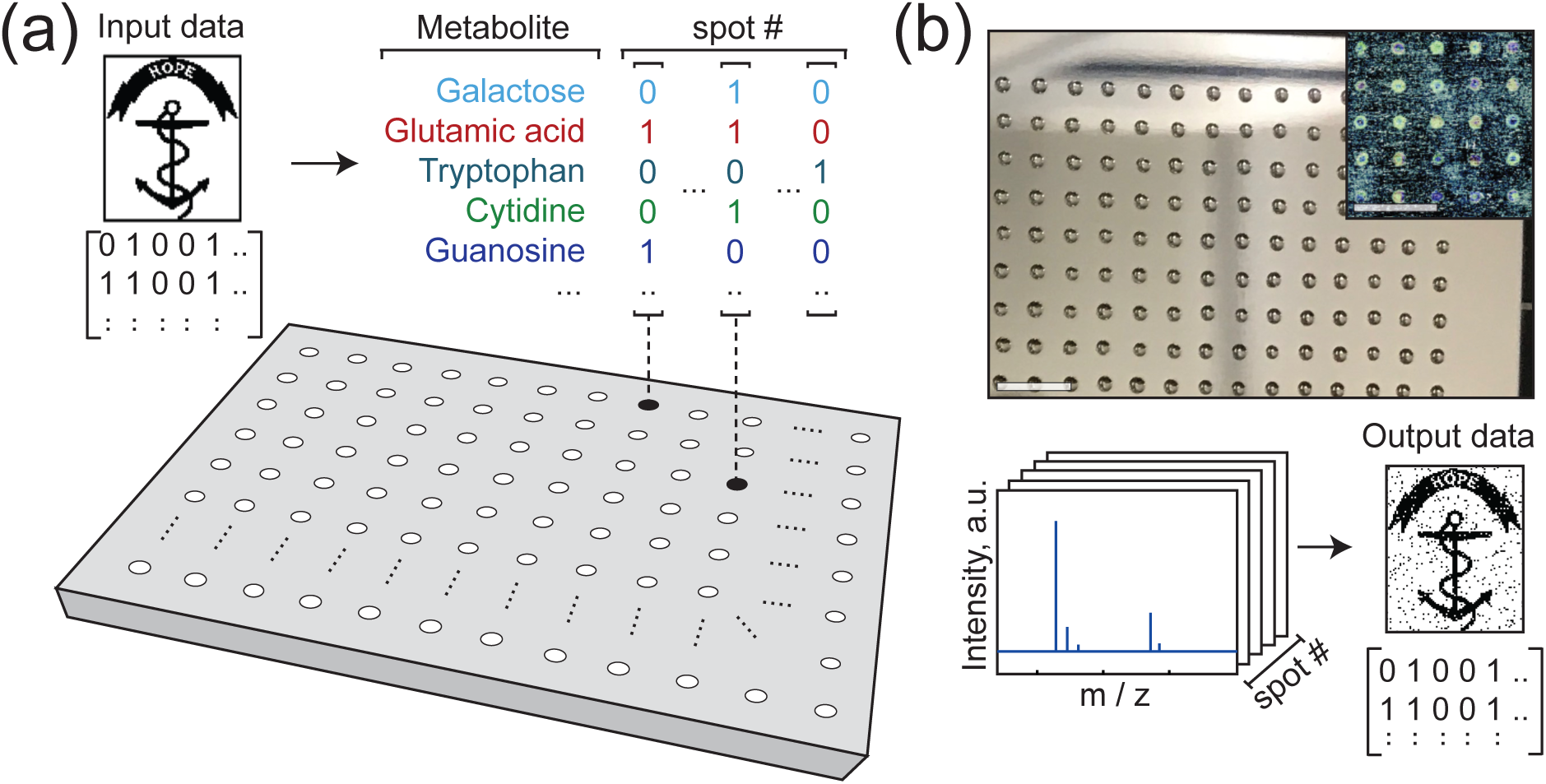
Writing and reading data encoded in mixtures of metabolites. **(a)** Binary image data is mapped onto a set of metabolite mixtures, with each bit determining the presence/absence of one compound in one mixture. For example, a spot mapped to four bits with values [0 1 0 1] may contain the 2^*nd*^ and 4^*th*^ metabolite at that location. (b) Small volumes of the mixtures are spotted onto a steel plate and the solvent is evaporated (scale bars: 5 mm). This chemical dataset is analyzed by MALDI mass spectrometry (b, bottom). Using the observed mass spectrum peaks, decisions are made about which metabolites are present. These decisions are assembled from the array of spots to recover the original image. The image shown is the Rhode Island Hope Regiment Colors [28].

We recover the encoded data from metabolic mixtures using mass spectrometry (Fig. 1b). The data aquisition is inherently parallelized, because a single mass spectrum provides information on every compound in a mixture. Noise characterization and logistic regression strategies for recovering the data are presented, along with examples of chemically encoded digital images. Raw error rates <1% are achieved with kilobyte-scale data sets using a simple peak analysis, illustrating the viability of both writing and reading metabolomic information. We use these experimental demonstrations to consider the benefits and limitations of encoding data into a biochemical medium in which interactions and interconversions can occur.

## Results

### Writing Synthetic Metabolomes

Our synthetic metabolome is a diverse set of 36 compounds (see Supplementary Table 1) including vitamins, nucleosides, nucleotides, amino acids, sugars, and metabolic pathway intermediates (all purchased from Sigma-Aldrich). To write data with the synthetic metabolome, we use an acoustic liquid handler (Echo 550, Labcyte) to transfer pure metabolic solutions in 2.5 nL increments to pre-defined locations on a steel MALDI plate. This produces a spatial array of different mixtures of metabolites (see Methods) where the presence (or absence) of each compound in each mixture encodes one bit of information.

After evaporating the solvent, each data plate contains up to 1536 dried spots on a 2.25 mm pitch (Fig. 1b), which we can analyze using Matrix Assisted Laser Desorption Ionization (MALDI) mass spectrometry (MS). To prescreen each compound in the synthetic metabolome, a plate was written with combinatorial mixtures of all 36 metabolites across 1400 unique spots (see Supplemental Figure S1). Since MALDI protocols are chemically specific, we do not expect the same identification accuracy across the whole compound library under one set of conditions. We use this pre-screen to determine the MS identification accuracy for every metabolite with the same protocol.

### Ion Cyclotron Mass Spectrometry of Metabolite Mixtures

A Fourier-transform ion cyclotron resonance (FT-ICR) mass spectrometer (SolariX 7T, Bruker) was used to analyze the array of crystallized mixtures. In FT-ICR MS, a pulsed RF field excites ions into a periodic orbit with a frequency that is given by the magnetic field strength and the ion’s mass [29], which enables much finer mass resolution than time-of-flight (ToF) instruments. In these experiments, the mass resolution is typically 0.001 Da (see Supplemental Figure S2). Using FT-ICR MS, metabolites can be discriminated even if their masses are only milli-Daltons apart.

In Fig. 2(a), one positive-ion MALDI-FT-ICR mass spectrum is shown for a spot which included guanosine (*go*), together with 9-Aminoacridine (*9A*) matrix. Protonated matrix adducts are identified at peaks 1 and 6 (blue), along with adducts of guanosine, labeled (2: Na, 3: K, 4: 2K – H and 5: isoproyl alcohol (IPA) + H). The observed intensities vary by adduct and species. In Fig. 2(b), the intensity of the first peak (protonated matrix at m/z = 195.0916±0.001) is illustrated across 1024 spots.

**Figure 2:**
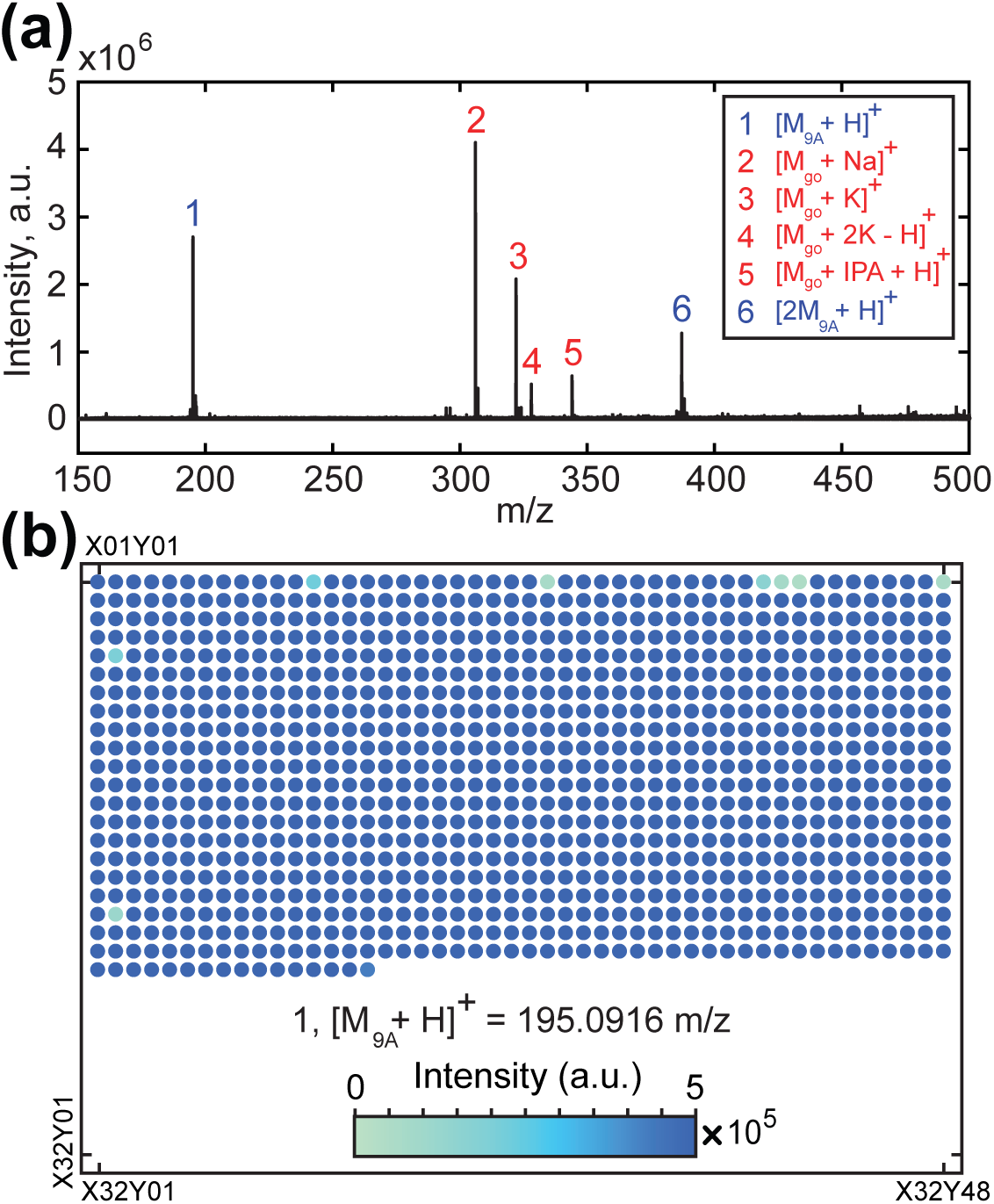
Analyzing chemical data plates with mass spectrometry. **(a)** Positive mode MALDI-FT-ICR mass spectrum of one spot containing Guanosine (*go*) and the MALDI matrix, 9-Aminoacridine (*9A*). Automated analysis of each spot used 4x averaged 1-second acquisitions. *go* ions (2, 3, 4, 5, in red) are present, along with two protonated matrix peaks (1, 6, in blue). **(b)** The intensity of the protonated matrix (peak 1) at m/z = 195.0916 ± 0.001 is shown graphically for a MALDI plate with 1024 independent mixture spots. Protonated aminoacridine is positively identified in 1020 spots (99.6 %).

Many open-access tools are available for metabolite peak detection and assignment from MS spectra [21]. To clearly relate the mass spectra to binary data, we consider a rudimentary detection scheme: if a metabolite’s mass intensity is above a particular threshold, then it is declared present, and the binary state of its address is set to 1 (or to 0, if its mass peak is absent). This approach identified the substrate matrix protonated peak in 1020 out of 1024 spots (≈99.6 %) in Fig. 2(b).

As an inital demonstration, we selected a library subset of 6 metabolites, which were used to encode a 6,142-pixel binary image of a Nubian ibex [30] into an array of 1024 mixtures (see Supplementary Figure S3). After pseudo-random interleaving, the data was mapped onto the presence or absence of sorbitol (*so*), glutamic acid (*ga*), tryptophan (*tp*), cytidine (*cd)*, guanosine (*go*) and 2-deoxyguanosine hydrate (*gh*). The plate was written and then analyzed using FT-ICR MS as detailed in the Methods.

Fig. 3a presents a spatial map and histogram of the spectral background noise observed in 240 independent spots. Before further analysis, we divide each spectrum by its background *σ*, which allows more direct comparison of signal strength at multiple locations. Signal strength is a complex function of the sample preparation, analyte and adduct. After normalization, peaks of interest for the 6 metabolites are shown in Fig. 3b. The first row is a spot whose data contains the six bits [1 0 0 0 0 0], and thus only the m/z peak associated with the first metabolite (sorbitol) is present. Similarly, five other ‘one-shot’ patterns are shown which can be decoded without error.

**Figure 3:**
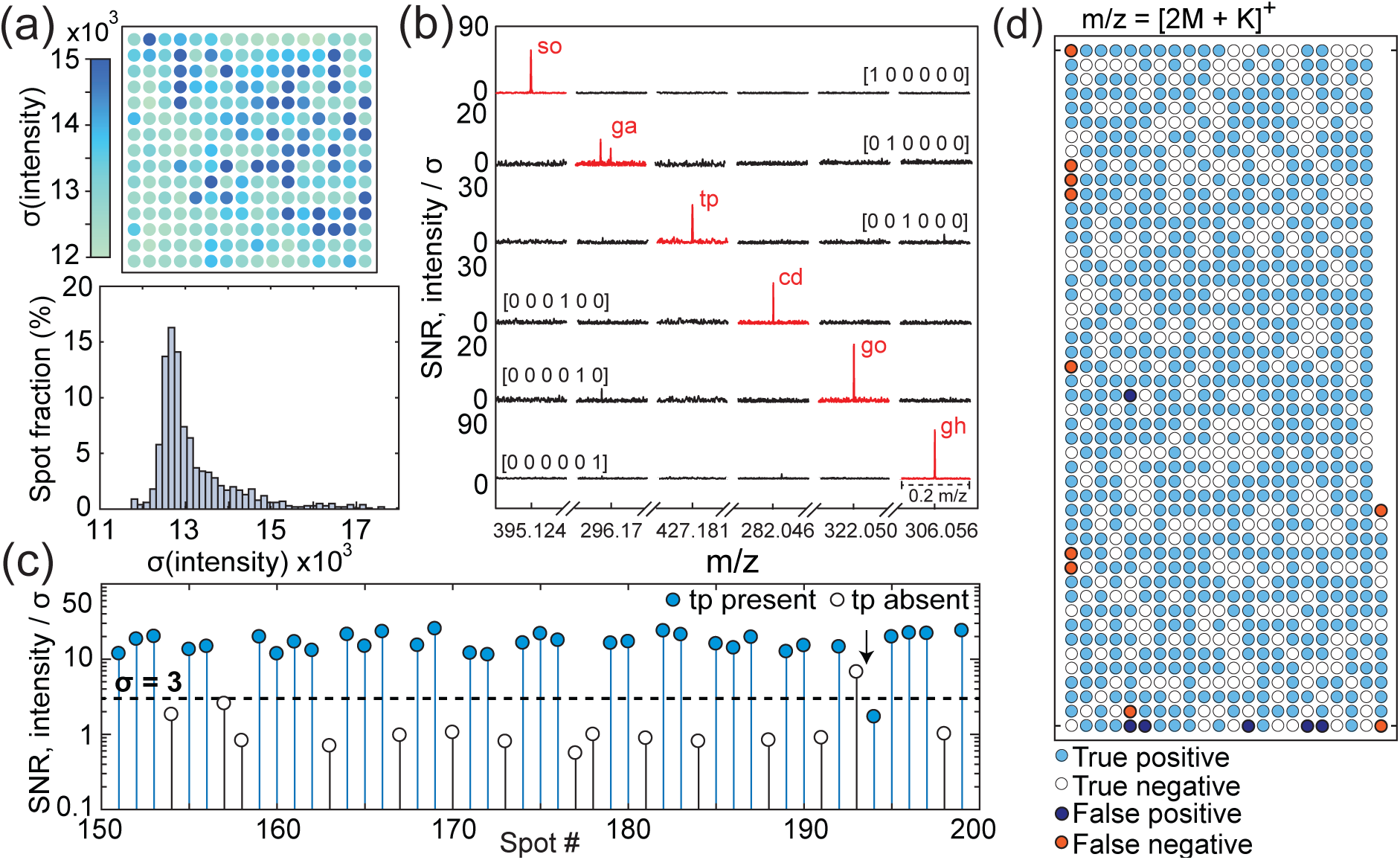
Spectral background and noise considerations. **(a)** Heatmap of the standard deviation of the MALDI-FT-ICR-MS spectral background noise from 240 unique spots of distinct mixtures, and a histogram of the background intensity standard deviation. **(b)** Spectra for six metabolites, normalized by the noise standard deviation. The m/z is cropped to six ranges of interest. Six ‘one-hit’ mixtures are plotted, one for each metabolite. **(c)** To assign presence/absence, we choose an intensity threshold at an appropriate m/z. As shown here, a 3*σ* threshold applied to the [2M_*tp*_ + K]^+^ tryptophan peak yielded a discrimination accuracy of 96%. **(d)** A hit map of the same tryptophan peak illustrating recovery using the 3*σ* threshold. Interestingly, the few errors are clustered at the edges of the plate.

A threshold of 3*σ* was chosen as the intensity required to declare the presence of a metabolite. For example, if we examine the tryptophan [2M_*tp*_ + K]^+^ mass (Fig. 3c), we find that this threshold yields 96% correct classification. This detection scheme can also be visualized for each spot on the plate, as shown in Fig. 3d. Clustering of errors at the edges of the plate suggests that small misalignments between the MALDI laser positions and the droplet spotting locations were a source of error.

### Statistical Analysis of Data Plates

In practice, one compound will be associated with multiple peaks, having varying signal-to-noise ratios and usefulness. For a given metabolome, we should attempt to identify which m/z peaks are most appropriate to identify each library element. Each high-resolution FT-ICR mass spectrum contains ∼2 × 10^6^ m/z points. Since most of the spectral space is background, it is helpful to first reduce the number of features to those which may be statistically useful. 1,444 candidate peaks found in the ensemble average of all mass spectra were tested to determine how accurately the intensities at that m/z classified the encoded data values (Fig. 4a).

**Figure 4:**
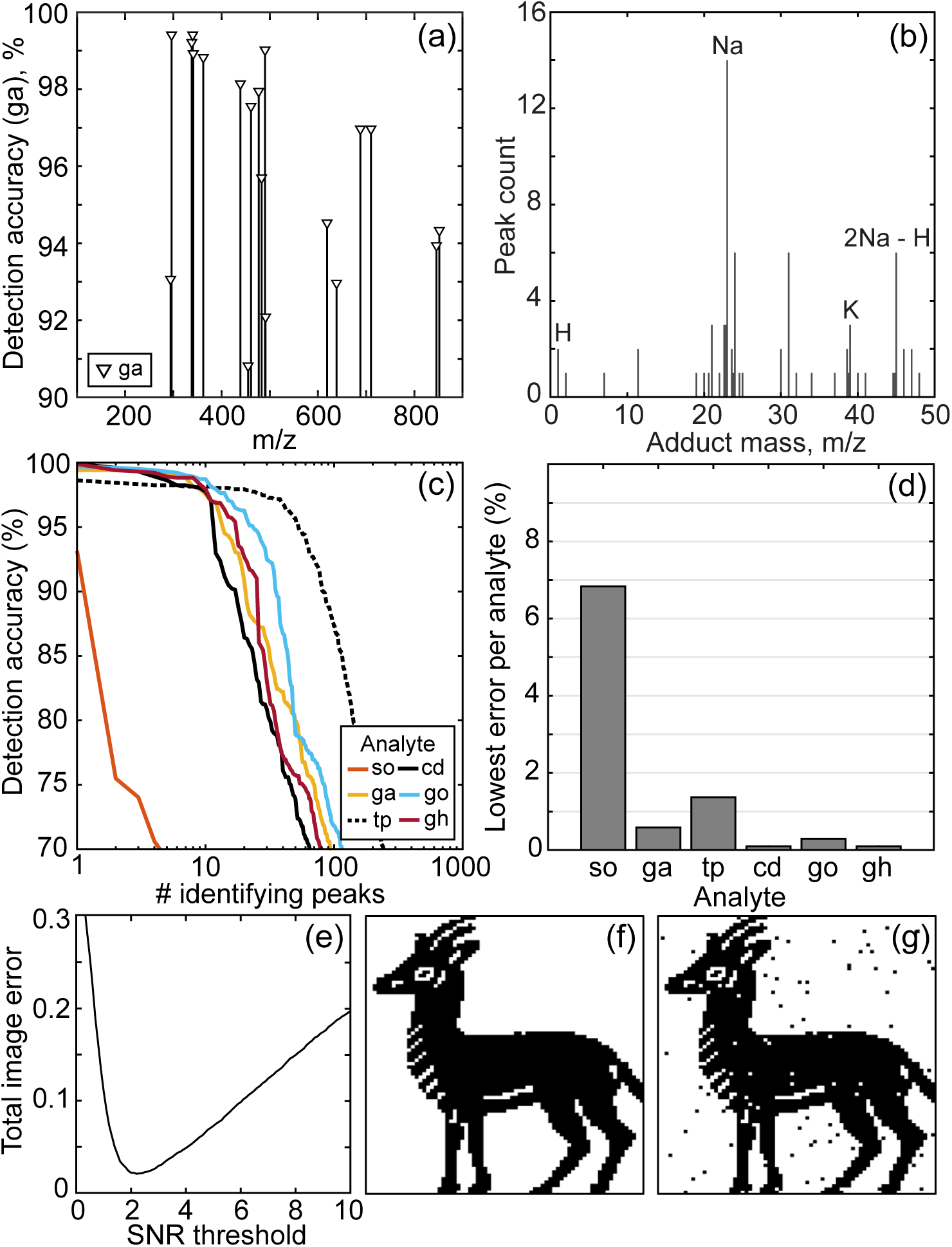
Identifying discriminating peaks. **(a)** The read recovery for different masses across the MS spectrum is shown for *ga*. **(b)** The histogram of adducts associated with peaks from the data in Fig. 3 indicates sodiated ions are predominant. **(c)** For each metabolite, we plot the number of peaks achieving a given detection accuracy. With the exception of sorbitol, each metabolite has at least 10 identifying peaks with >97% accuracy. **(d)** The error of the single best performing mass for each metabolite. **(e)** Using only the best performing mass from (d), the error rate for the six metabolites across 1024 locations (6144 bits) is shown as a function of the SNR cutoff. These mixtures encoded the 6142-bit image shown in **(f)**. In **(g)**, we recover the image with a 2.5*σ* decision threshold, producing approximately 2% cumulative read/write error.

Although these peaks were identified without chemical bias, many features can be attributed to known metabolite adduct ions (although some are synthesis byproducts or derive from the substrate matrix). A histogram of the associated adduct masses is shown in Fig. 4b. H, Na, Na-H and K adducts are all frequently observed.

The number of peaks achieving detection accuracy in the range of 70-100% is shown in Fig. 4c. Selecting the best performing peak for each metabolite, and applying a detection threshold of 2.5*σ*, was sufficient to recover data at about 2% cumulative read/write error (Fig. 4e). The corresponding input and output data images are shown in Fig. 4f-g. The simplicity and success of the overall read and write process is encouraging, but there is still significant room for improvement.

### Decoding Data from Multiple Peaks using Logistic Regression

Assuming that the discriminating peaks are partially uncorrelated (see Supplemental Figure S4), it is reasonable to seek improvement by utilizing multiple m/z peaks per metabolite. Such strategies will become increasingly important in more complex metabolomes.

Using techniques similar to those for the 6kb ibex image, we encoded a 17,424-bit image of a cat from an Egyptian tomb [31] using 1,452 spots containing data mixtures from a 12-metabolite subset of the library (Fig. 5a). We used this data to extend the decoding scheme to incorporate multiple m/z features. After identifying the set of statistically discriminating peaks, we performed a logistic regression using between 1 and 16 of the best-performing peaks. Multi-mass linear regression achieved a read accuracy of 97.7% for the whole cat image (Fig. 5c). Cumulative read error rates for the data in Fig. 4 and Fig. 5 are shown as a function of the number of masses used in the logistic regression. Applying these techniques to the earlier ibex dataset, an error rate of <0.5% was achieved. However, repeated measurement of spots can cause data loss. It was found that <1% error was added by each successive read of a data plate (see Supplementary Figure S5). Using different plates, one for training the regression, and one recovering data, achieved the same error rates and no overfitting (see Supplementary Figure S6). Overall, these demonstrations show that the metabolome is a viable and robust medium for representing digital information.

**Figure 5:**
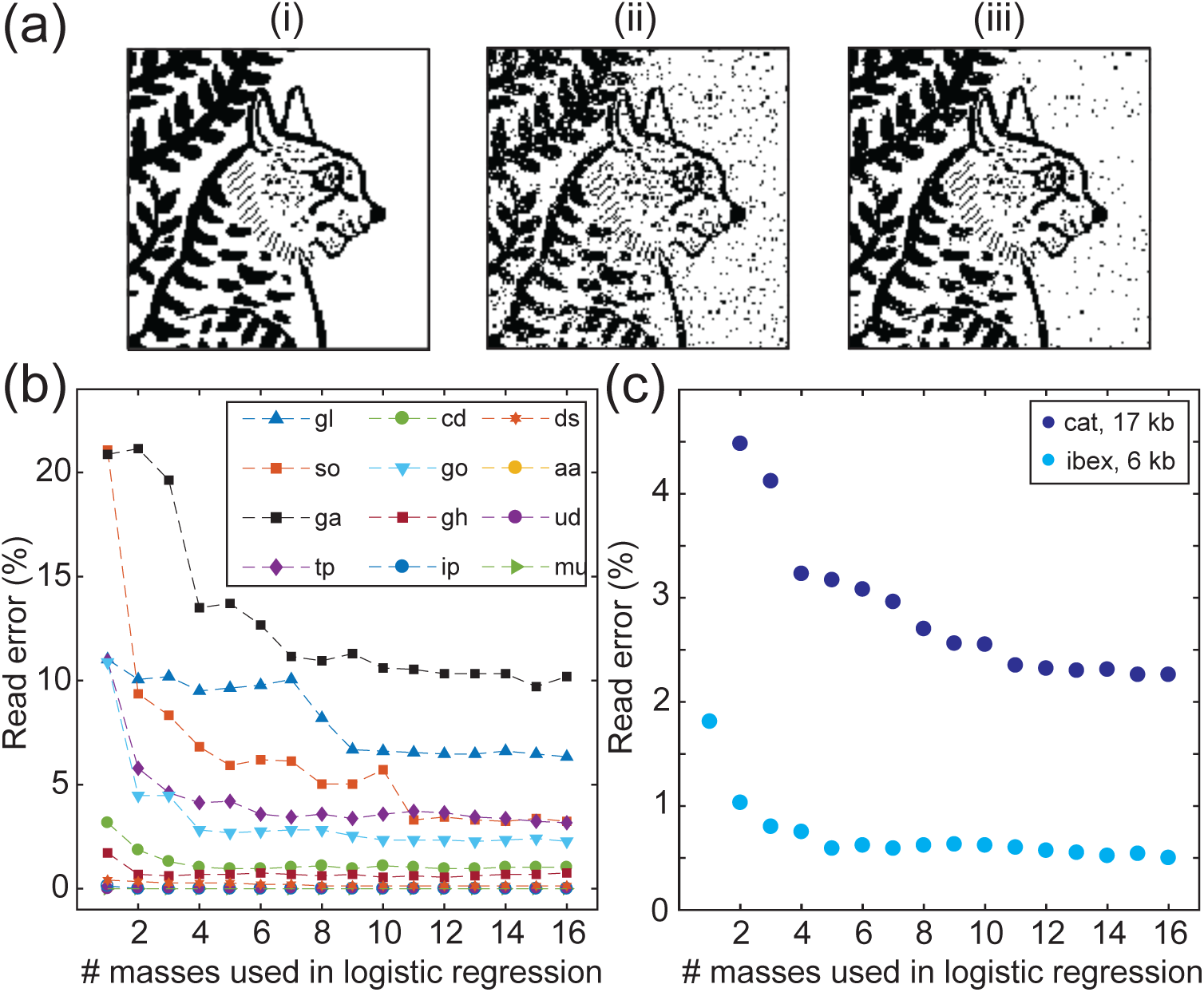
Logistic regression for multi-peak molecular data readout. Improvements over single-peak classification can be achieved with logistic regression utilizing multiple identifying masses per metabolite. **(a)** (i) A 17,424-bit image written into 1452 mixtures from a 12-metabolite subset of the library. (ii) The image recovered using one discriminating mass per metabolite. (iii) The image recovered using a regression combining 16 peaks per molecule. **(b)** Some compounds achieve low error rates even with single peaks. However, other molecules do not have an isolated m/z peak that shows high accuracy by itself. For these compounds, multi-peak logistic regression improves classification. (c) Cumulative read error rates for the two images as a function of the number of masses used in the logistic regression.

## Discussion

One advantage of molecular data storage is its high storage density. To date, demonstrations using DNA have reached about 214 petabytes per gram [32], although this is still orders of magnitude from theoretical limits [33]. An encoded metabolome written using a large small-molecule library could improve on this number [34], but our experiments highlight several limitations and potential benefits that warrant further discussion.

Statistically discriminating m/z features were used to classify the metabolite mixtures and recover the data at 98-99.5% accuracy using a simple analysis. Further development can take advantage of the wide range of sophisticated analysis technologies for metabolic profiling, including artificial neural networks, genetic algorithms, and self-organizing maps [35]. The inclusion of these methods, in conjunction with error correcting codes, leaves ample headroom for improved data recovery from more complex mixtures.

In terms of data rates, we demonstrated write speeds of 5 bits/sec, and aggregate read speeds of 11 bits/sec. We have performed little optimization of either the read or write times, and as the size of the metabolite library is increased, the MS read speed in particular has significant room to improve.

Looking forward, it is interesting to consider the upper bound on information capacity using all known metabolites (∼10^5^ [14]). Even if only a fraction are stable, detectable, and display unique masses, this conservatively predicts hundreds of bits per spectral acquisition, which could all be read in parallel. As sub-zeptomole MS and nanomolar concentration detection have been available for nearly two decades [36, 37], detection at this level of complexity seems plausible.

Improvements in spatial density, and perhaps write speed, could come from reducing the volume and pitch of spots. There are opportunities for high density multilayer printing. To avoid storage density limits arising from finite transfer volumes, the precise mixture of metabolites associated with one spot can be pre-mixed in one well of an intermediary data plate. Transfer of 2.5 nL from the intermediary plate well to one spot means that hundreds of metabolites can be present in a nL volume on the plate. There is also room to extend on this work using larger libraries for higher capacity, or by storing multiple bits per complex, leveraging oligomerization [38].

As a proof of principle we chose nL spot sizes for ease of handling and convenience. Scaling the mixture spots down to diffraction-limited laser spot scales (from 2.25 mm to ∼ *µ*m spot pitch) would improve data storage density by 6 orders of magnitude. Theoretically, this could facilitate extension from multi-kilobyte to multi-gigabyte data sets per plate. However, the true limit of data storage density depends on the availabley instrumentation.

ICR-MS (or other high-resolution MS such as orbital traps) have a finite ion capacity per acquisition, so the number of compounds can not be arbitrarily increased due to competition. Metabolites with a lower ionization efficiency will be excluded even though present in a large, competitive mixture. Therefore, to increase the number of metabolites per spot, future work may need to screen libraries for ionization efficiency. Alternatively, it may be that other read strategies (e.g. nanopores [39,40]) would provide higher sensitivity than MS, but the protocol demonstrated here has ample room to increase capacity by several orders of magnitude.

A likely source of error in more complex mixtures will be interactions between metabolites [5]. However, interspecies networks may also have benefits, such as opportunities for overwriting or transforming data, which hints at possibilities for synthetic metabolomic computation. One recurring challenge in metabolomics is obtaining trustworthy ‘ground truth’ samples. Perhaps by considering metabolomes as more abstract and mutable stores of information, we can develop new tools that allow us to overcome statistical biases, establish ground truths, and tease out subtle interactions and interconversion rates in well-regulated synthetic metabolomes.

## Conclusion

‘Omics’ technologies have grown out of genomics to encompass other complex information-rich systems like the metabolome. It is only natural to ask whether there exist complementary opportunities to make use of metabolites’ structural diversity and interactivity. As a proof of principle of postgenomic information storage, we have experimentally encoded >100,000 bits of digital images into synthetic metabolomes (see Supplementary Table 2), and we are confident that this number can be increased significantly in the future. One novel contribution is the demonstration of data storage in a mixture of dissimilar molecules, which can improve information capacity and read times through diversity and parallelism. Perhaps more importantly, this work offers a new perspective on chemical information, and it introduces possibilities for synthetic metabolomic computation and establishing metabolic ‘ground truths’ through interrogation of synthetic metabolomes.

## Methods

### Chemical Preparation

Reagent grade samples of 36 distinct metabolic compounds (see Supplementary Table 1) were diluted in dimethyl sulfoxide (DMSO, anhydrous), each to a nominal concentration of 25mM. Some metabolites were initially dissolved in an alternative solvent (de-ionized water with or without 0.5M or 1M hydrochloric acid) to facilitate solvation in DMSO. 10*µL* of each compound was aliquoted into a 384-well microplate (Labcyte 384LDV).

### Data Mixture Preparation

The chemical mixtures were prepared on a 76 × 120 mm^2^ stainless steel MALDI plate. An acoustic liquid handler (Labcyte Echo 550) was employed to transfer the compounds from the library wellplate onto the MALDI plate. The nominal droplet transfer volume is 2.5 nL, but to reduce variability, we typically use 2 droplets (5 nL) per compound. The destinations of the droplets are programmed to match a standard 2.25mm pitch 1536-spot (32 × 48) target.

After spotting the compounds to the MALDI plate, a MALDI matrix material was added to each location. We selected 9-Aminoacridine for its compatibility with metabolite libraries, its low background in the small molecule regime, and its support for both positive and negative ion modes. The MALDI plate is left to dry and crystallize overnight (∼ 10 hours). Once dried, the plate can be stored in a humidity controlled cabinet or analyzed by MALDI-FT-ICR mass spectrometry.

### Mass Analysis of Data Plates

A Fourier-transform ion cyclotron resonance (FT-ICR) mass spectrometer (SolariX 7T, Bruker) was used to analyze the crystallized metabolite data mixtures. The exact resolution is a function of the measurement time allocated per spectrum. For these experiments, we typically used 0.5-1 sec, yielding a resolution of < 0.001 Da. The instrument is run in MALDI mode and is configured to serially measure the mass spectrum of each mixture on the 48×32 grid. Acquisiton for a full plate takes <2 hours.

In multinomial logistic regression, the probability of a metabolite being present is modeled as a combination of multiple predictor masses. Detection considers the natural exponent of an offset plus the sum of all identifying mass SNRs, where each SNR is multiplied by a trained weight coefficient. A limited-memory BFGS algorithm was used to predict the multinomial logistic accuracy scores given an input of the *n* best peaks per metabolite. This process was iterated for all metabolome constituents.

## Acknowledgements

The authors are grateful for support from Sherief Reda, Eunsuk Kim, and Jason Sello. This research was supported by funding from the Defense Advanced Research Projects Agency (DARPA W911NF-18-2-0031). The views, opinions and/or findings expressed are those of the authors and should not be interpreted as representing the official views or policies of the Department of Defense or the U.S. Government.

## Supplementary Information

### Synthetic metabolome library

**Table 1:** A list of screened compounds. The shown mass is the monoisotopic mass, as found on PubChem [1].

| Key | Name | Description | CID | Mass |
| --- | --- | --- | --- | --- |
| **aa** | cis-Aconitic acid | acid | 643757 | 174.016 |
| **cm** | Creatine monohydrate | ADP to ATP recycler | 80116 | 149.08 |
| **es** | Sodium ethyl sulfate | alcohol metabolite | 23680278 | 147.981 |
| **gm** | Glutamine | amino acid | 738 | 146.069 |
| **mt** | Methionine | amino acid | 6137 | 149.051 |
| **hd** | Histidine | amino acid | 6274 | 155.069 |
| **pa** | Phenylalanine | amino acid | 6140 | 165.079 |
| **la** | L-(+)-Arginine | amino acid | 6322 | 174.112 |
| **ts** | Tyrosine | amino acid | 6057 | 181.074 |
| **tp** | Tryptophan | amino acid | 6305 | 204.09 |
| **ga** | N-Acetyl-L-glutamic acid | arginine intermediate | 70914 | 189.064 |
| **dr** | D-(-)-Ribose | carbohydrate | 5311110 | 150.053 |
| **ip** | Imidazolepropionic acid | histidine metabolite | 70630 | 140.059 |
| **pp** | 4-Hydroxyphenylpyruvic acid | keto acid | 979 | 180.042 |
| **ad** | Adenosine 5'-diphosphate sodium | energy recycler | 6022 | 427.029 |
| **gl** | D-(+)-Galactose | monosaccharide | 6036 | 180.063 |
| **td** | Thymidine | nucleoside | 5789 | 242.09 |
| **cd** | Cytidine | nucleoside | 6175 | 243.086 |
| **ud** | Uridine | nucleoside | 6029 | 244.07 |
| **da** | 2'-Deoxyadenosine | nucleoside | 13730 | 251.102 |
| **mu** | 5-Methyluridine | nucleoside | 445408 | 258.085 |
| **as** | Adenosine | nucleoside | 60961 | 267.097 |
| **dm** | 2'-Deoxyadenosine monohydrate | nucleoside | 9549172 | 269.112 |
| **go** | Guanosine | nucleoside | 6802 | 283.092 |
| **gh** | 2'-Deoxyguanosine hydrate | nucleoside | 16218597 | 285.107 |
| **ct** | Cytidine 5'-triphosphate NaCl | nucleoside | 16219171 | 526.948 |
| **gp** | D-Glucose 6-phosphate sodium salt | pathway initiator | 23702133 | 282.012 |
| **na** | NADP Disodium | pathway initiator | 2734411 | 787.039 |
| **dc** | 2'-Deoxycytidine | pyrimidine metab. | 13711 | 227.091 |
| **du** | 2'-Deoxyuridine | pyrimidine metab. | 13712 | 228.075 |
| **ds** | Disodium succinate | salted acid | 9020 | 161.99 |
| **sc** | Trisodium Citrate Dihydrate | salted acid | 71474 | 293.994 |
| **so** | Sorbitol | sugar alcohol | 5780 | 182.079 |
| **th** | Thiamine hydrochloride | vitamin | 6202 | 336.058 |
| **rf** | Roboflavin | vitamin | 493570 | 376.138 |
| **pq** | Phylloquinone | vitamin | 5284607 | 450.35 |

**Figure 1:**
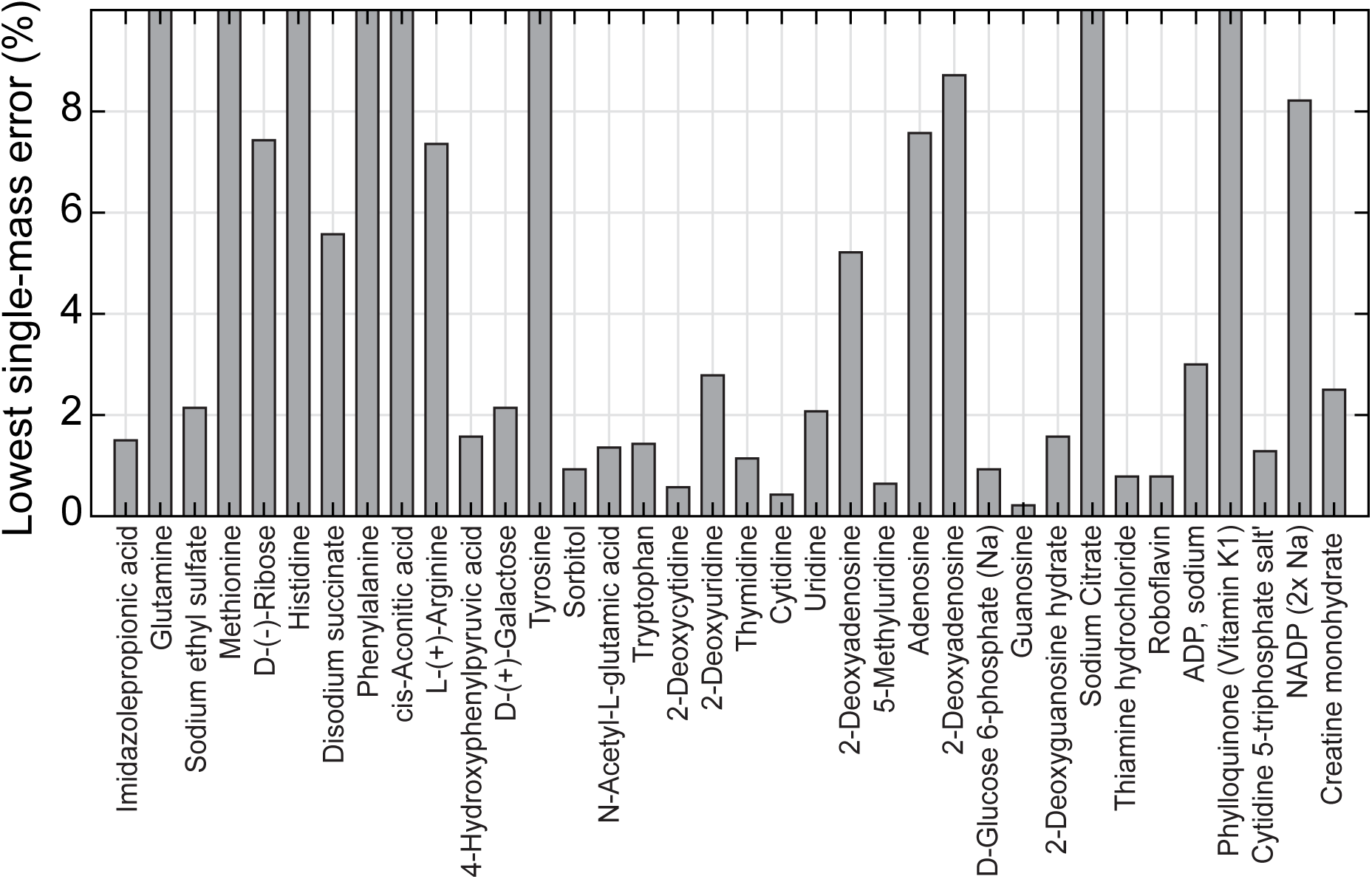
Read error rates for each component of the synthetic metabolome. The data is derived from a 1400-spot plate, where each metabolite was prescribed pseudo-randomly as present or absent. Since each spot contained a mixture of 3 present and 33 absent metabolites, the error rates shown consider a degree of mixture error due to metabolic conversion. 8 / 36 metabolites have single-best-peak error rates > 10%, possibly due to poor uptake and solvation in DMSO. About half of the compounds yielded single-best-peak error rates of < 2%.

**Table 2:** A list of image data sets written, with the number of repetitions. Cumulatively, ≈ 108,700 bits were written into synthetic metabolomes.

| Description | Ref. | bits | repetitions | net kbits |
| --- | --- | --- | --- | --- |
| Ibex print | [2] | 6,142 | 2× | 12.3 |
| RI Flag 1781 | [3] | 8,904 | 3× | 26.7 |
| Cat drawing | [4] | 17,424 | 4× | 69.7 |

**Figure 2:**
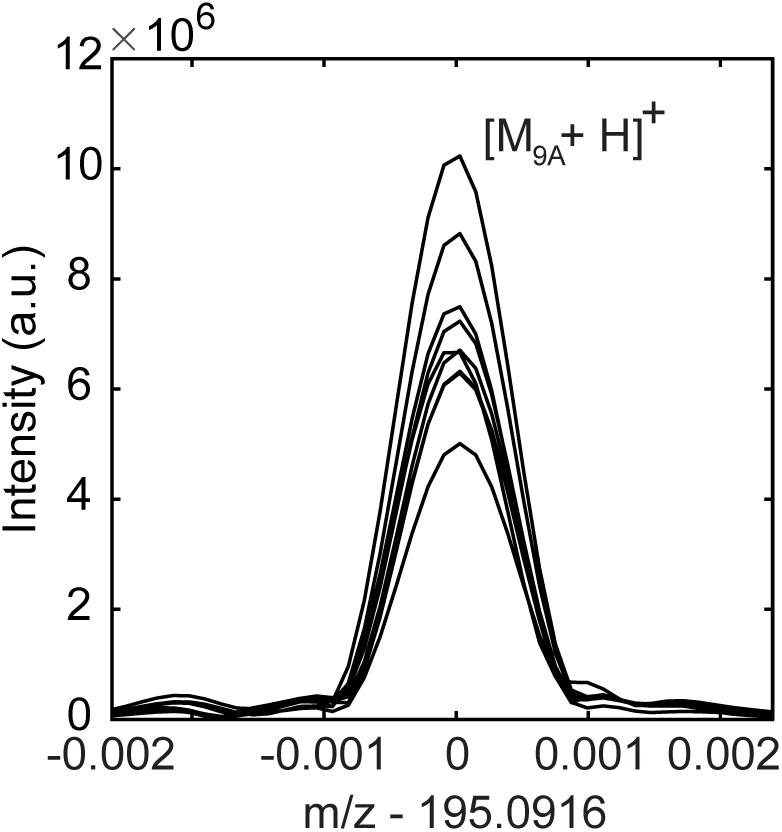
Eight repetitions of MALDI mass spectral acquisition are shown centered at m/z = 195.0916 (protonated 9-Aminoacridine). Each repetition is from a unique deposition of 40 nL of 18.25 mM matrix in DMSO (air dried). The entirety of the peak above background is captured within the spectral window range M ± 0.001 m/z, regardless of signal intensity.

**Figure 3:**
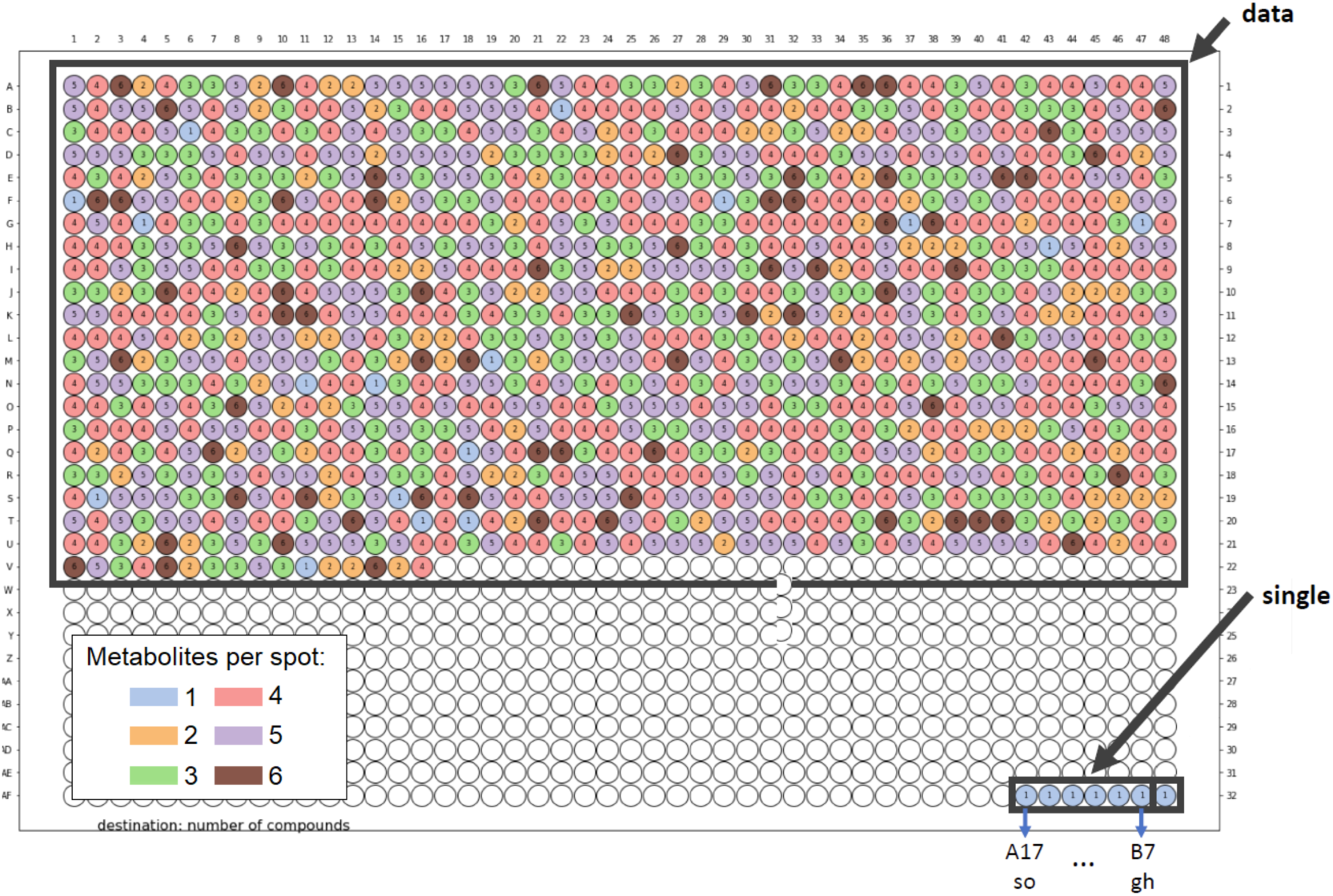
(a) A graphical overview of the Nubian ibex data plate contents. A color-coding is used to denote the number of metabolites present in each of the 1024 spot mixtures.

**Figure 4:**
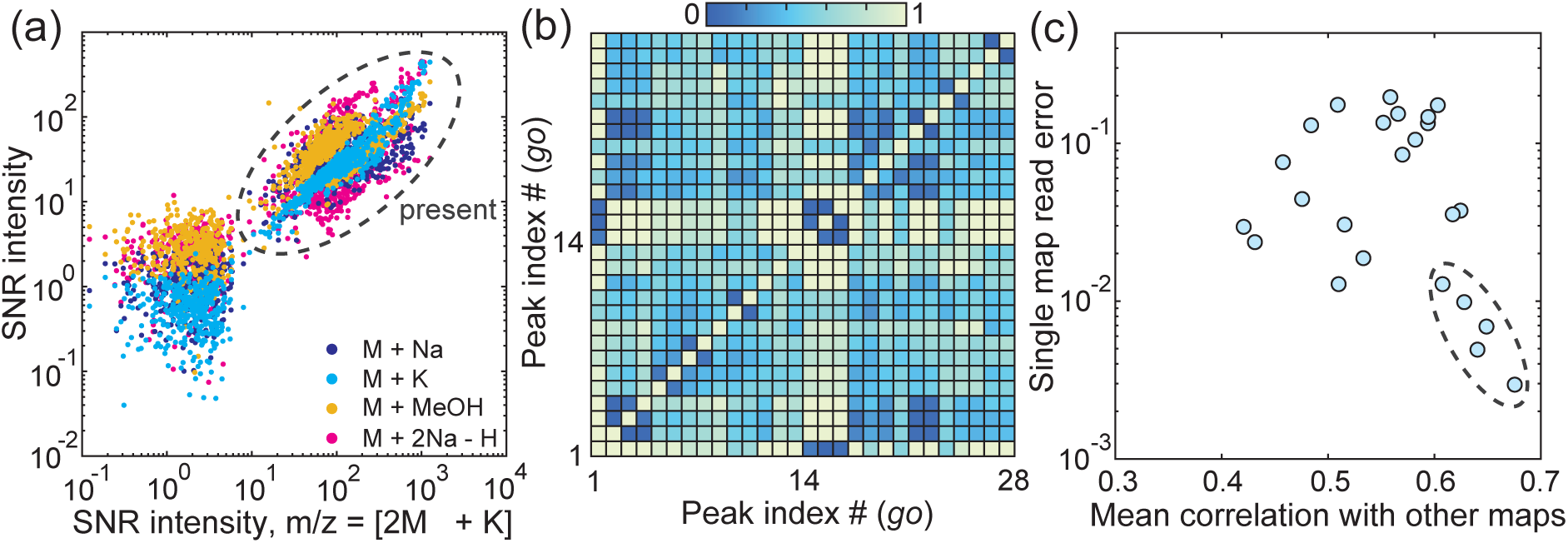
Appreciating the degree of correlation in readout errors using multiple m/z peaks. Each mass is associated with a list of normalized intensities from each spot on a plate. (a) The normalized intensity of the [2M+K]^+^ peak from all 1024 locations is shown for guanosine, plotted against the intensities of other guanosine ions. The intensities clustered into present (dotted ellipse) and absent states, and ion intensities are positively but imperfectly correlated. In (b), the 28 best discriminating masses are selected for autocorrelation. Some sets of masses exhibit clustered groupings, but correlations are imperfect. (c) The effective read error at each m/z is plotted against its mean correlation with other guanosine features. The masses which yield the lowest errors are often more correlated (dotted ellipse).

**Figure 5:**
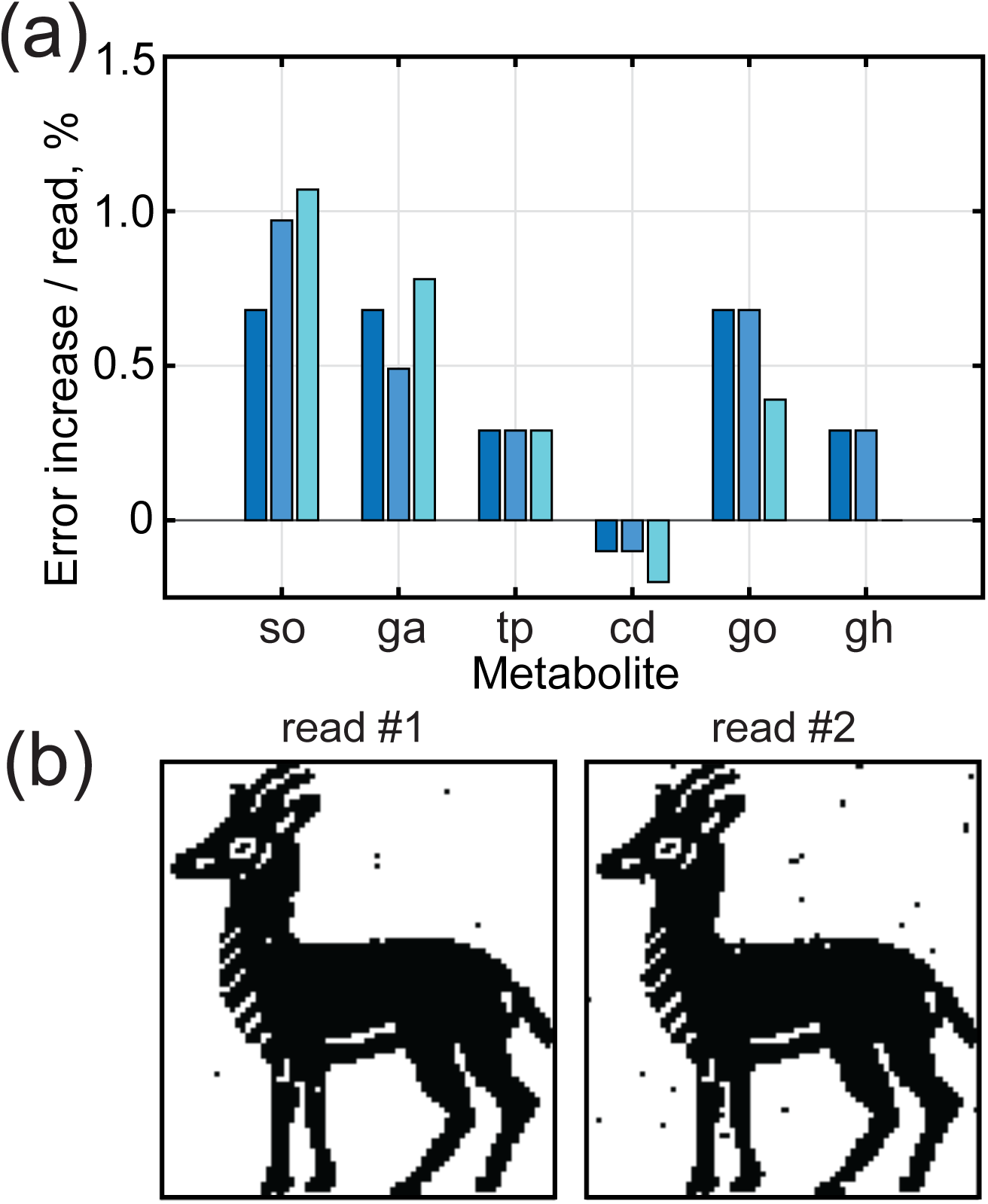
Repeated measurement of spots can cause data loss. A 6142 bit image data plate was written using 6 metabolites as described in Figure 4. The plate was read several times. The increase in error rate per read is shown in (a) broken out by metabolite. The first and second read repetition of the image using 16-peak logistic regression (see Methods) are shown in (b). Each read took <2 hrs. Typically, <1% error was added by each successive measurement of a data plate.

**Figure 6:**
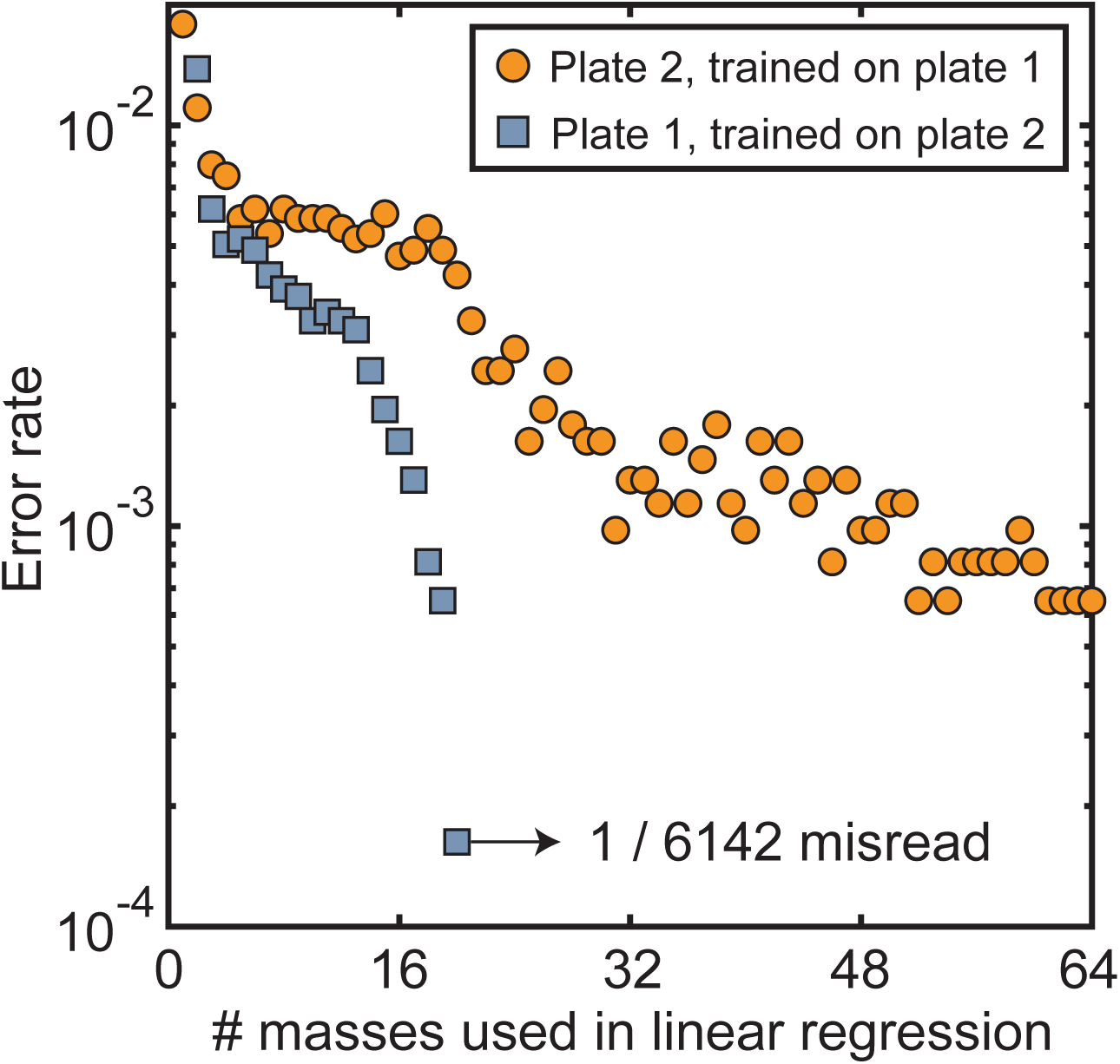
Cross validation. Two identical 6142 bit image data plates, plate 1 and plate 2, were both written using the 6 metabolites described in Figure 4. Plate 1 was used to train the logistic regression to determine which masses were optimal discriminators. These masses were then used to recover data from plate 2. The process was then reversed. Error rates are shown as a function of the number of identifying masses used in regression, where training and testing of the data has been seperated as described. Although some plate-specific complexity is evident, aggregate error rates dropped below 0.1% in both cases, with complete seperation of training and test data.

**Table 3:**
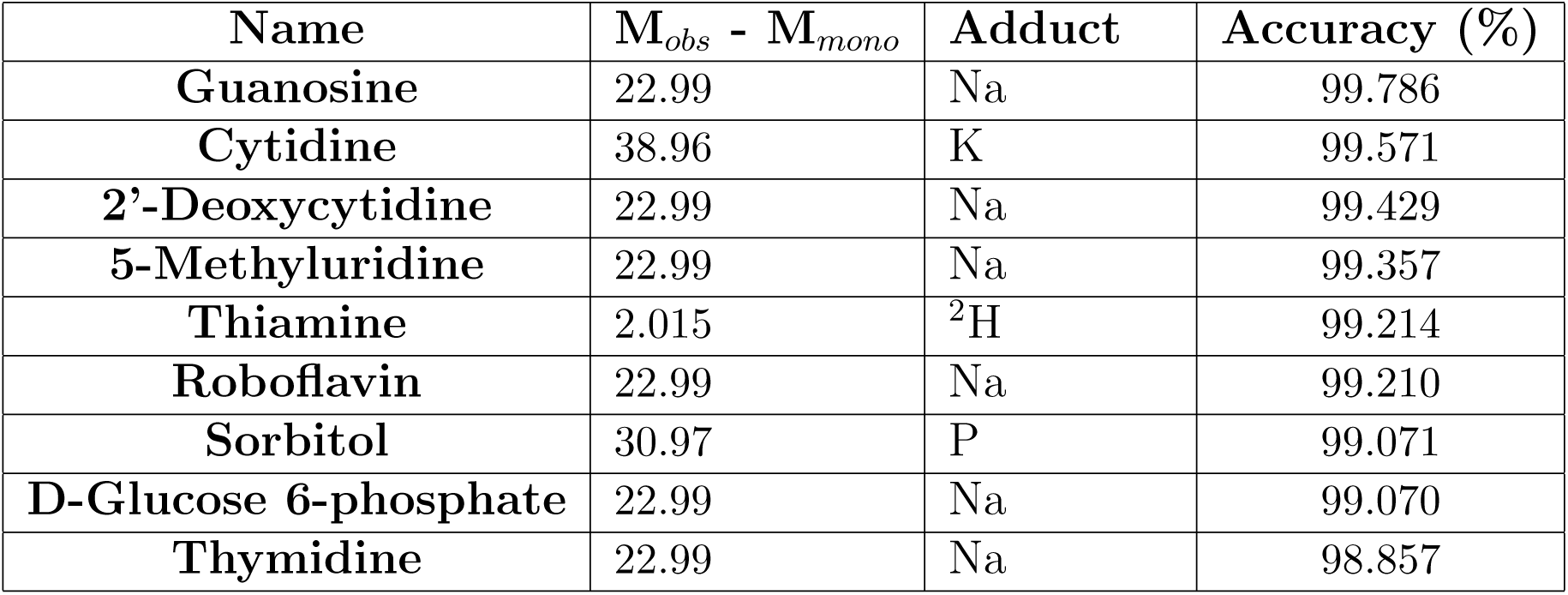
Adduct classification and data recovery accuracy. The best discrminating peak mass minus the monoisotoic mass (M_*obs*_ – M_*mono*_) is shown for metabolites. The adduct type is determined from the residual mass. The accuracy recovered from each of the adducts is shown.

| Name | $M_{obs} - M_{mono}$ | Adduct | Accuracy (%) |
| --- | --- | --- | --- |
| Guanosine | 22.99 | Na | 99.786 |
| Cytidine | 38.96 | K | 99.571 |
| 2'-Deoxycytidine | 22.99 | Na | 99.429 |
| 5-Methyluridine | 22.99 | Na | 99.357 |
| Thiamine | 2.015 | $^2\text{H}$ | 99.214 |
| Roboflavin | 22.99 | Na | 99.210 |
| Sorbitol | 30.97 | P | 99.071 |
| D-Glucose 6-phosphate | 22.99 | Na | 99.070 |
| Thymidine | 22.99 | Na | 98.857 |

**Table 4:** Guanosine adduct list. The discrminating peak mass minus the monoisoptic mass (M_*obs*_ – M_*mono*_) is shown for 10 peaks associated with Guanosine. The adduct type is determined from the residual mass. The data recovery accuracy for each mass multiple and adduct is shown. For large absolute m/z, the error in m/z increases due to finite sampling limitations.

| Multiple (n) | $M_{obs} - n \times M_{mono}$ | Adduct | Accuracy (%) |
| --- | --- | --- | --- |
| 1 | 22.99 | Na | 99.786 |
| 1 | 23.99 | Na + $1_c$ | 98.926 |
| 1 | 38.96 | K | 99.604 |
| 1 | 44.97 | 2Na - H | 98.729 |
| 2 | 22.99 | Na | 95.2150 |
| 2 | 38.95 | K | 94.7270 |
| 2 | 23.99 | Na + $1_c$ | 94.0430 |
| 2 | 30.97 | P | 91.1300 |
| 2 | 44.96 | 2Na - H | 90.7230 |
| 2 | 39.95 | K + $1_c$ | 88.6700 |

